# β-catenin-dependent endomesoderm specification appears to be a Bilateria-specific co-option

**DOI:** 10.1101/2022.10.15.512282

**Authors:** Tatiana Lebedeva, Johan Boström, David Mörsdorf, Isabell Niedermoser, Evgeny Genikhovich, Igor Adameyko, Grigory Genikhovich

## Abstract

Endomesoderm specification based on a maternal β-catenin signal and axial patterning by interpreting a gradient of zygotic Wnt/β-catenin signalling was suggested to predate the split between Bilateria and their evolutionary sister Cnidaria. However, in Cnidaria, the roles of β-catenin signalling in both these processes have not been proven directly. Here, by tagging the endogenous β-catenin protein in the sea anemone *Nematostella vectensis*, we show that the oral-aboral axis in a cnidarian is indeed patterned by a gradient of β-catenin signalling. Unexpectedly, in a striking contrast to Bilateria, *Nematostella* endoderm specification takes place opposite to the part of the embryo, where β-catenin is translocated into the nuclei. This suggests that β-catenin-dependent endomesoderm specification is a Bilateria-specific co-option, which may have linked endomesoderm specification with the subsequent posterior-anterior patterning.

## Main text

During the early development of bilaterian embryos, β-catenin signalling is involved in two fundamental processes occurring sequentially: it specifies the endomesoderm, and it patterns the posterior-anterior (P-A) axis (*1-11*). The central role of the Wnt/ β-catenin (cWnt) signalling in the patterning of the main body axis is not restricted to Bilateria. Expression data indicate that cWnt signalling may regulate axial patterning in the earliest branching metazoan groups Ctenophora and Porifera (*12-14*), and functional analyses of the last 25 years showed the involvement of the graded cWnt signalling in the patterning of the oral-aboral (O-A) axis in the bilaterian sister group Cnidaria (sea anemones, corals, jellyfish, *Hydra*) (*15-21*). Moreover, nuclear localization of β-catenin on one side of the sea anemone embryo at the early blastula stage and the failure to gastrulate upon β-catenin loss-of-function suggested that the involvement of β-catenin signalling in the endomesoderm specification was also an ancestral feature conserved at least since before the cnidarian-bilaterian split over 700 Mya (*17, 22-24*). However, in spite of convincing circumstantial evidence, there was no direct proof for either the presence of the O-A gradient of β-catenin signalling or for the instructive role of β-catenin signalling in the endomesoderm specification in a cnidarian. Here we ventured to obtain such proof and close this knowledge gap by tagging endogenous β-catenin in a model cnidarian – the sea anemone *Nematostella vectensis* – with superfolder GFP (*25*) and detecting its localization at the time of germ layer specification and in the axial patterning phase.

Previously we showed that genes expressed in distinct ectodermal domains along the O-A axis in *Nematostella* react dose-dependently to different levels of pharmacological upregulation of the β-catenin signalling (*17*). Downstream, β-catenin signalling activates a set of transcription factors, among which the more orally expressed ones act as transcriptional repressors of the more aborally expressed ones (*16*). Like this, the initially ubiquitous aboral identity of the embryo is restricted in a β-catenin-dependent manner to the future aboral domain as the oral and the midbody domains appear and become spatially resolved. In this process, JNK signalling appears to act agonistically with β-catenin signalling: JNK inhibitor treatment aboralizes the embryo, and JNK inhibition is also capable of dose-dependently rescuing the oralization caused by pharmacological upregulation of β-catenin signalling with a GSK3β inhibitor (Supplementary Fig. 1). The regulatory logic of the β-catenin-dependent axial patterning and the complement of the downstream transcription factors is so strikingly similar between the sea anemone and the deuterostome bilaterians that it suggests the homology of the cnidarian O-A and the bilaterian P-A axis (*1-3, 16, 26, 27*).

To directly verify the presence of the oral-to-aboral β-catenin signalling gradient and the role of β-catenin signalling in the *Nematostella* endoderm specification, we used CRISPR/Cas9-mediated genome editing to generate a knock-in line, in which the nucleotides coding for the first 5 amino acids of β-catenin were replaced by the superfolder GFP (sfGFP) coding sequence. In order to test for the presence of the nuclear β-catenin gradient along the oral-aboral axis of the embryo, we incrossed heterozygous F1 polyps (*wild type/sfGFP-β-catenin*) and allowed the offspring to develop until late gastrula stage. As expected, approximately 3/4 of the embryos were fluorescent, however, fluorescent microscopy of live embryos only revealed strong signal at the cell boundaries – in line with the function of β-catenin in the cell contacts. In order to detect nuclear sfGFP-β-catenin, we fixed the embryos and stained them with an anti-GFP antibody. Antibody staining revealed a comparatively weak but clear nuclear signal forming an oral-to-aboral gradient in the ectoderm (Fig. 1A).

**Figure 1.**
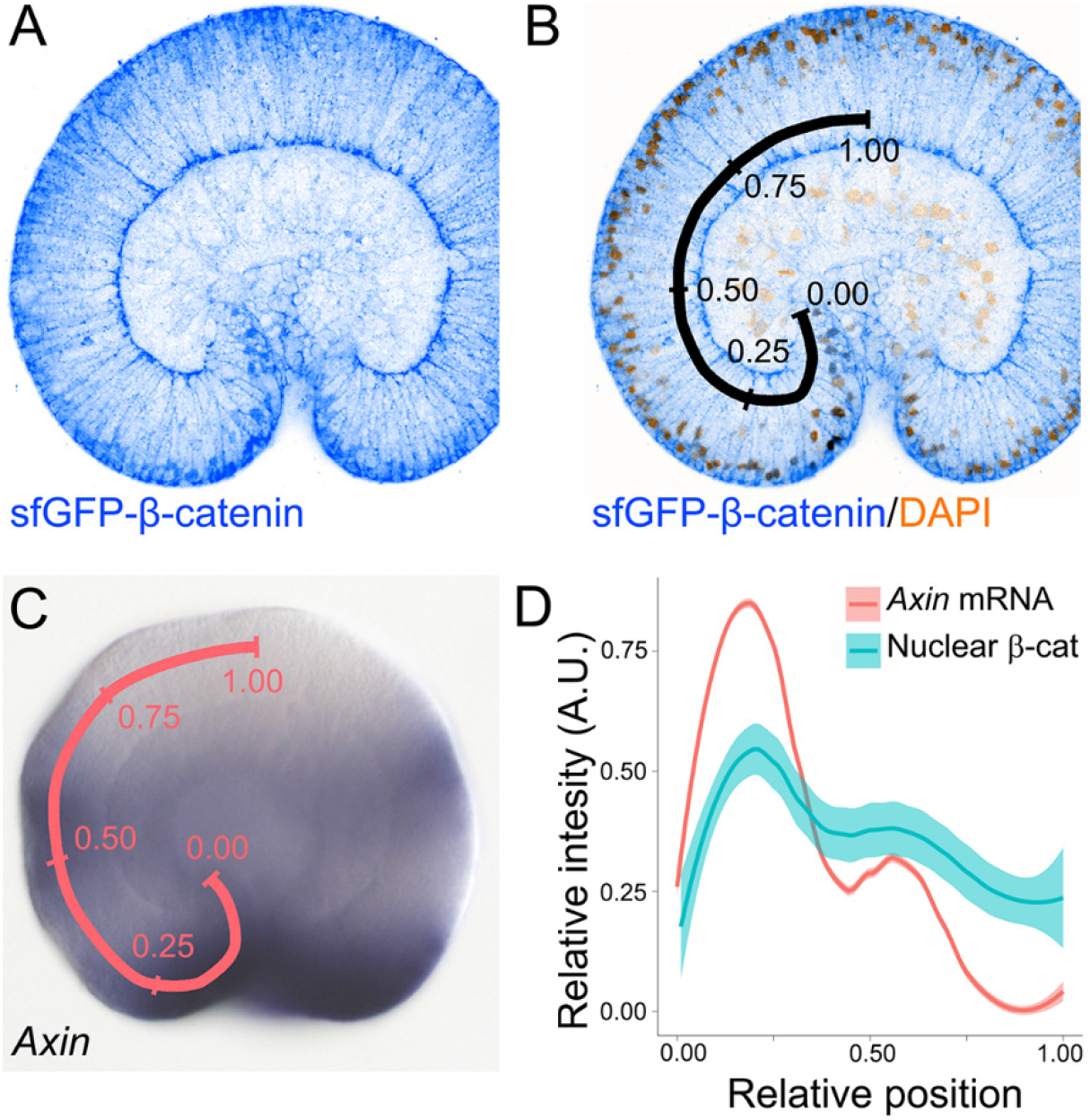
Nuclear sfGFP-β-catenin forms a bimodal oral-to-aboral gradient in late gastrula stage embryos. (A) Anti-GFP antibody staining detects sfGFP-β-catenin at the cell membranes and in the nuclei. (B) Overlay of the anti-GFP signal with the nuclear staining shows the positions, at which anti-GFP staining was quantified. The first nucleus, where anti-GFP staining intensity was measured is located at the relative position 0.00; the last nucleus – at the relative position 1.00. (C) *Axin* in situ hybridization staining intensity was measured along the pink line from the relative position 0.00 to the relative position 1.00. (D) LOESS smoothed curves show that nuclear sfGFP-β-catenin forms an oral-to-aboral gradient with two peaks (n=6). These peaks correspond to the positions where β-catenin target *Axin* expression peaks as well (n=10).

Quantification of the signal intensity in all ectodermal nuclei starting from the deepest cell of the pharyngeal ectoderm (relative position 0.00) and ending with the cell in the centre of the aboral ectoderm (relative position 1.00) showed a peak of nuclear sfGFP-β-catenin in the bend of the blastopore lip (approximately at relative position 0.20), and a second, smaller peak at the border between the midbody and the aboral domain (approximately at relative position 0.60). Both peaks coincide with the peaks of expression of the conserved and highly sensitive β-catenin signalling target *Axin* (*15-17*) (Fig. 1B-D). Thus, we conclude that the initial assumption that genes expressed in distinct domains along the O-A axis and regulating its patterning react to a graded β-catenin signal is correct.

Our next goal was to verify the involvement of β-catenin signalling in the specification of the endoderm in *Nematostella*. Previous analyses of the *β-catenin-GFP* mRNA injected *Nematostella* embryos showed nuclear β-catenin-GFP localization on one side of the early blastula in the untreated embryos and in all blastoderm cells of the embryos upon GSK3β inhibition (*17, 22*). Moreover, *Nematostella* embryos with the nuclear localization of β-catenin suppressed either by truncated cadherin mRNA overexpression, injection of the β-catenin translation blocking morpholino or β-catenin RNAi did not form preendodermal plates, blastopore lips and failed to gastrulate remaining perfect blastula-like spheres (*22, 23, 30*). Morphologically, this effect perfectly resembled the gastrulation block caused by injection of the mRNA encoding truncated cadherin or the DIX domain of Dishevelled in sea urchin (*9*) and led to the conclusion that the endoderm in *Nematostella*, just like the endomesoderm in a number of bilaterians, is specified by an early β-catenin signal at the future gastrulation pole of the embryo (*22*). Although universally accepted in the field (also by us – see for example (*17, 31*)), this hypothesis was contradicted by several important observations (Fig. 2). First, in spite of lacking preendodermal plates, β-catenin morphants ubiquitously expressed endodermal marker *SnailA*, but not the zygotic aboral/anterior markers *Six3/6* and *FoxQ2a* (*16, 23*). Second, upon pharmacological activation of β-catenin signalling by GSK3β inhibitor treatment starting before 6 hours post-fertilization (hpf), the embryos also remained spherical failing to form preendodermal plates, blastopore lips and gastrulate; however, these spheres expressed oral ectoderm markers *Brachyury, FoxA* and *FoxB*, while endodermal markers were abolished (*15, 23*). Third, loss-of-function experiments showed that LRP5/6 and combined Fz knockdowns did not prevent endoderm specification and gastrulation in *Nematostella* in spite of completely aboralizing the ectoderm of embryo i.e. entirely disrupting its oral-aboral patterning (*15*). Taken together, these data suggest that, unlike the bilaterian endomesoderm, *Nematostella* endoderm specification seems to be repressed by β-catenin signalling rather than activated by it.

**Figure 2.**
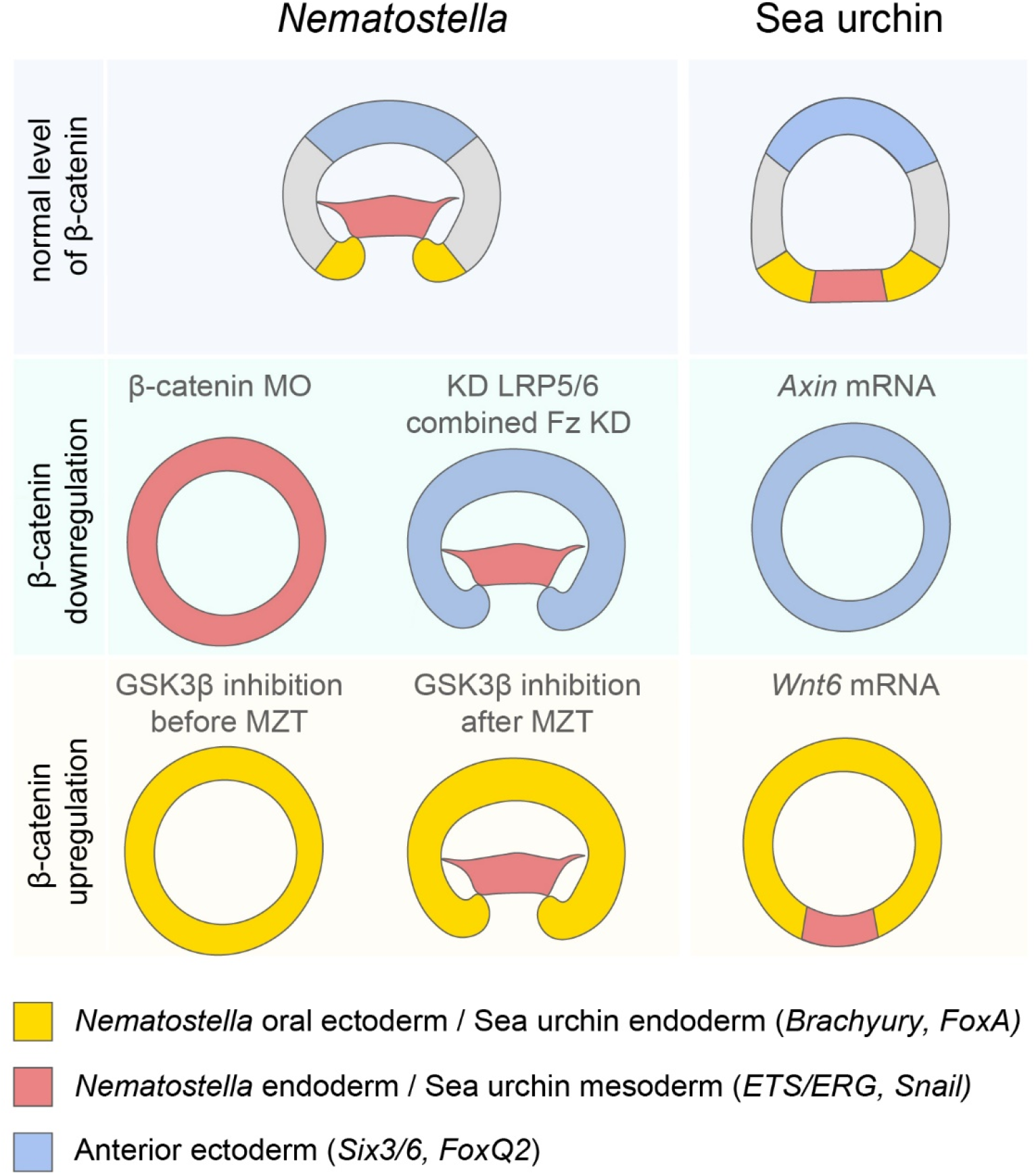
Comparison of the effects of the up- and downregulation of the β-catenin signalling in *Nematostella* and sea urchin. The germ layers are coloured according to the cnidarian-bilaterian germ layer equivalence hypothesis published in (*28*). The cartoons showing the effects of the up- and downregulation of the β-catenin signalling are based on data from (*15, 16, 23, 27, 29*). MZT – maternal-to-zygotic transition.

*Nematostella* endoderm specification is an early event happening at or prior to 6 hpf (*15, 16*), at which time nuclear sfGFP-β-catenin is clearly detectable in the developing embryos by fluorescent microscopy. Hence, in order to verify the involvement of β-catenin signalling in the specification of the endoderm, we immobilized sfGFP-β-catenin expressing embryos in low melting point agarose and performed live imaging from early cleavage until the onset of gastrulation (Fig. 3, Supplementary Movies 1-2). Nuclear sfGFP-β-catenin became detectable as early as during the 32-64 cell stage, and from the very start, nuclear signal was confined to approximately 2/3 of the embryo. Nuclear sfGFP-β-catenin became visible during every cell cycle until shortly after the desynchronization of the mitotic divisions at midblastula (*32*), after which it became too weak to detect, while the sfGFP-β-catenin signal in the cell contacts remained strong. Strikingly, in all embryos we live-imaged (n=10), the preendodermal plate formed on the side opposite to the side where nuclear sfGFP-β-catenin was detectable at earlier stages, i.e. early nuclear sfGFP-β-catenin was always observed on the aboral side of the embryo. In order to make sure that what we were observing was indeed the nuclear sfGFP-β-catenin dynamics, we also live-imaged sfGFP-β-catenin expressing embryos, which were incubated in the 5 μM solution of the GSK3β inhibitor alsterpaullone (ALP) from fertilization on (Supplementary Movies 3-4). In line with the previous publications (*17, 22*), upon GSK3β inhibition, fluorescent signal was observed in all nuclei from 16-32 cell stage on, and the embryos failed to gastrulate.

**Figure 3.**
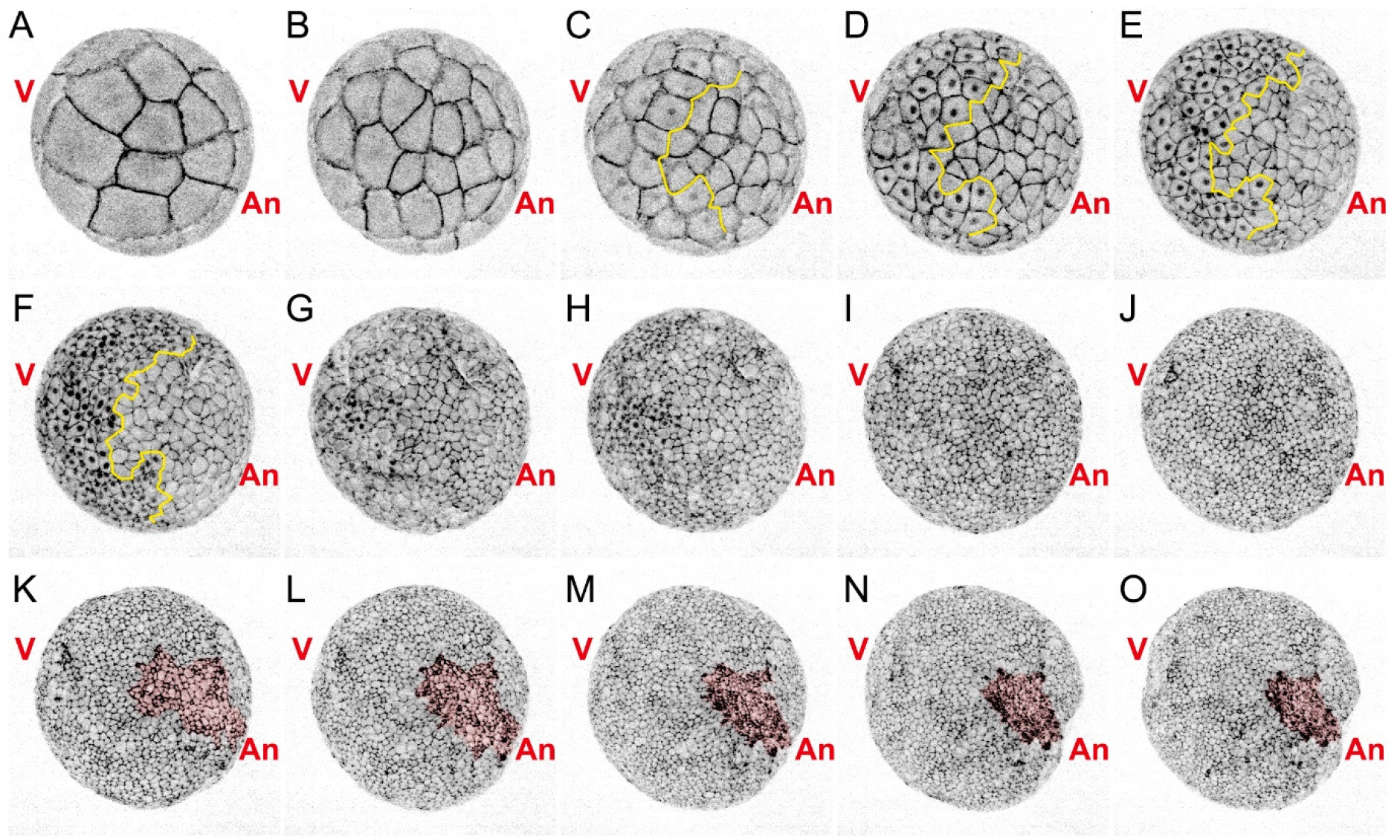
Early sfGFP-β-catenin accumulation is observed at the vegetal pole, opposite to the future gastrulation site. Individual frames from the Supplementary Movie 1 showing the same embryo over the course of development. sfGFP-β-catenin – black signal. An – animal pole, V – vegetal pole, the preendodermal plate is highlighted pink. Note nuclear sfGFP-β-catenin in the vegetal/aboral half of the embryo on (C-H). Yellow line on (C-D) demarcates the sharp boundary between the nuclear sfGFP-β-catenin-positive and nuclear sfGFP-β-catenin-negative cells until the loss of synchronicity in cell division on (G).

The presence of nuclear sfGFP-β-catenin on the aboral rather than oral side in the embryos developing in the absence of ALP (Fig. 3, Supplementary Movies 1-2) suggests that in *Nematostella*, unlike in Bilateria, instead of promoting endoderm specification, β-catenin signalling actually represses it, which resolves all the discrepancies mentioned above. First, it explains why endodermal marker *SnailA* is ubiquitously expressed in the β-catenin morphants but zygotic aboral/anterior ectoderm markers *Six3/6* and *FoxQ2a* are not (*23*). Second, it explains why upon treatment with a GSK3β inhibitor prior to 6 hpf, the whole embryo acquires the oral ectoderm fate rather than the endoderm fate (*15, 23*). Third, the lack of β-catenin signalling at the future oral side of the early embryo is in line with our observation that endoderm formation is not controlled by Fz/LRP5/6-mediated signalling (*15*).

Our finding that an oral-to-aboral gradient of nuclear β-catenin exists in the *Nematostella* gastrula confirms a number of previous assumptions on the mode and logic of the oral-aboral patterning in this animal and is in line with the idea that the cnidarian O-A axis corresponds to the bilaterian P-A axis (*15-17*). However, our second observation that *Nematostella* endoderm forms in the β-catenin-negative domain has a much greater import for the understanding of the early evolution of the body axes and germ layers. In Bilateria, unless physically prevented by large amounts of yolk, endomesoderm specification and gastrulation take place at the vegetal pole, i.e. posteriorly. In contrast, in Cnidaria, gastrulation modes are highly variable. Some species gastrulate by invagination, unipolar ingression or epiboly, while others have multipolar modes of gastrulation such as cellular, morular or mixed delamination or multipolar ingression (*33*). In case of multipolar gastrulation, germ layer specification and gastrulation movements are spatially uncoupled from the universally cWnt-dependent O-A patterning (*21, 34-37*). Importantly, in cnidarians with a unipolar mode of gastrulation, endoderm specification and gastrulation always takes place at the animal, rather than at the vegetal pole, and in all cnidarians analysed so far, the animal-vegetal axis exactly corresponds to the O-A axis of the embryo independent of whether they have a multipolar or a unipolar mode of gastrulation (*32, 33, 38-40*). Previously, it has been proposed that the activation of the β-catenin-dependent endomesoderm specification at the vegetal rather than at the animal pole of a stem bilaterian resulted in the inversion of the position of the gastrulation site in Bilateria (*41*). Our new data suggest a different scenario (Fig. 4). Both, in *Nematostella* and in Bilateria, maternal β-catenin accumulates in the vegetal pole nuclei, however, the specification of the endomesoderm by this signal appears to be a bilaterian specific co-option, which linked germ layer specification, gastrulation movements and P-A patterning. In contrast, in Cnidaria, endoderm specification appears to be either negatively controlled by β-catenin (as in *Nematostella*) or not to be controlled by β-catenin at all (as in cnidarians with multipolar gastrulation modes), which explains the variety of gastrulation modes observed in this phylum. Many new questions arise with this observation, for example: i) what causes endoderm specification and subsequent gastrulation movements in the β-catenin-negative domain in *Nematostella* and in random positions in cnidarians with multipolar gastrulation modes, or ii) what regulatory changes tethered bilaterian gastrulation to the ancestral site of the nuclear β-catenin accumulation at the vegetal pole? However, our data clearly suggest that the current view on the ancestral mode of endomesoderm specification in animals needs to be re-assessed.

**Figure 4.**
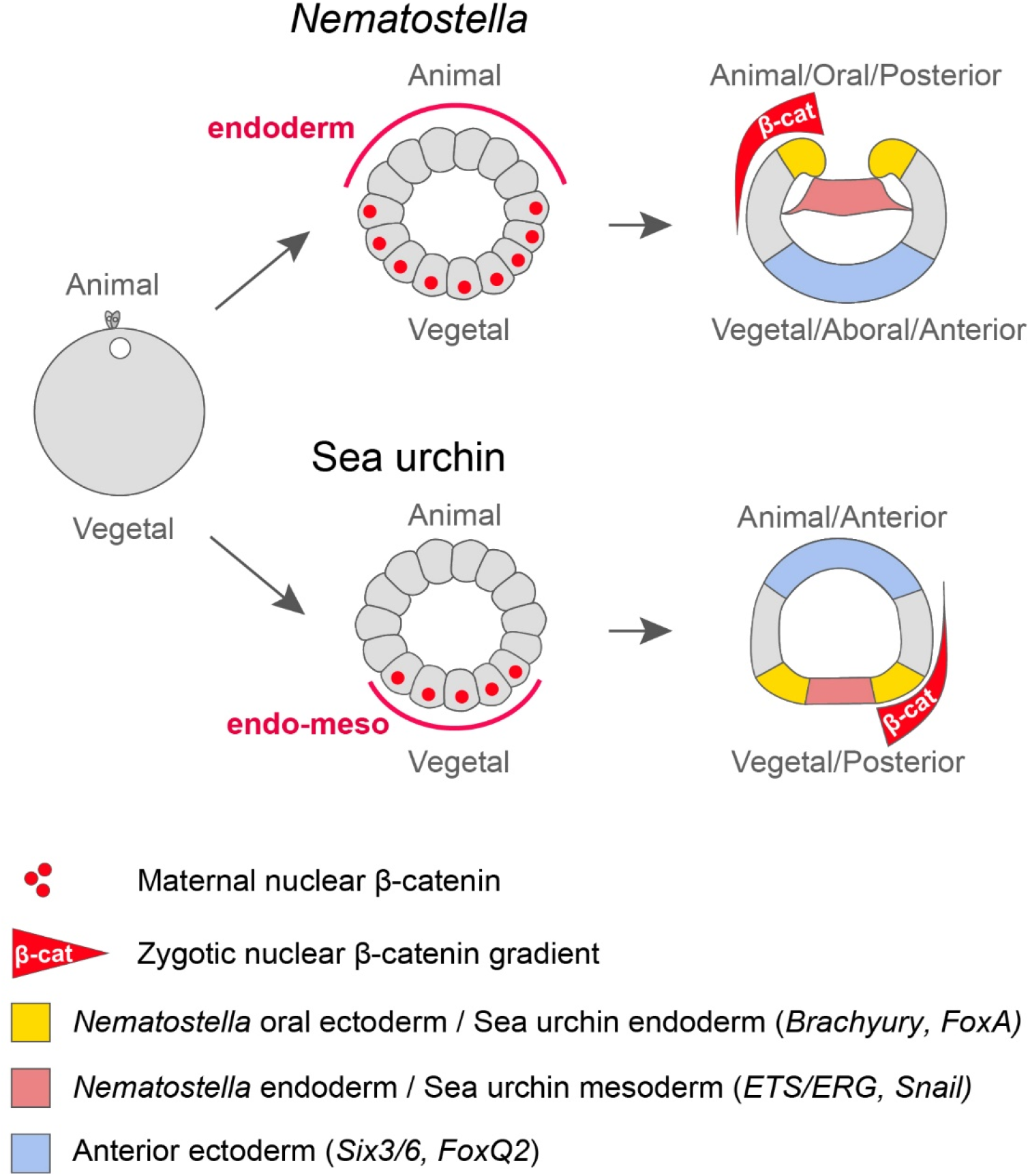
β-catenin signalling may have been co-opted for endomesoderm specification at the base of Bilateria.

## Materials and methods

### Animal culture and generation of the sfGFP-β-catenin knock-in line

The animals were maintained and spawning was induced as described in (*42*). To generate a *sfGFP-β-catenin* knock-in, a single gRNA 5’-ACCATGGAGACACACGGTAT-3’ recognising a sequence starting at the position 134602 on the scaffold_183 of the *Nematostella* genome v.1 (*43*) was selected using CHOPCHOP (*44*), and CRISPR/Cas9 genome editing was performed as described in (*45*). For homologous recombination, we generated a fragment in which the first five triplets of the *Nematostella β-catenin* coding sequence were replaced with the *Superfolder GFP* coding sequence introduced in frame by Gibson assembly. The fragment containing the homology arms and *sfGFP* was amplified using the primers 5’-GTGGAATTCGCAGCATTTCTCA-3’ and 5’-TCAAGGATGGCTCAGCAAGC-3’, which were modified as described in (*46*). F0 animals with clear fluorescent patches were raised to sexual maturity and crossed to wild type to generate heterozygous F1. To confirm the knock-in, we clipped single tentacles from individual heterozygous F1 animals, extracted genomic DNA from them and performed PCR using the primers 5’-GGTCGTAGATGGTACCCTAAG-3’ and 5’-CAACTCTGGGATAGCACGTGTAG-3’ located in the *β-catenin* genomic locus upstream and downstream of the homology arms. This PCR resulted in two *β-catenin* locus fragments with and without the *sfGFP* insertion, which we confirmed by Sanger sequencing. Genotyped knock-in animals were raised to maturity, sexed and intercrossed. The offspring of these genotyped F1 animals was used in the experiments.

### Antibody staining, in situ hybridization and staining intensity measurements

For anti-GFP antibody staining, the embryos were fixed for 1 hr in 4%PFA/PBS-TT (PBS-TT = 1x PBS containing 0.2% Tween20 and 0.2% TritonX100) at 4°C, washed five times for 5 min in PBS-TT, incubated for 2 hours in a blocking solution consisting of 95% BSA/PBS-TT and 5% heat inactivated sheep serum (BSA/PBS-TT = 1% BSA w/v in PBS-TT), and stained overnight at 4°C in rabbit polyclonal anti-GFP (abcam290) diluted 1:500 in the blocking solution. Unbound antibody was removed by five 15 min washes in PBS-TT, then the embryos were blocked again and stained overnight at 4°C with AlexaFluor488 donkey anti-rabbit IgG (Life Technologies) diluted 1:1000 in the blocking solution. The unbound secondary antibody was removed by 5 washes with PBS-TT; DAPI was added to the first wash to counterstain the nuclei, then the embryos were gradually embedded in Vectashield (Vectorlabs). 16 bit images of the DAPI and anti-GFP staining were obtained using the Leica SP8 LSCM equipped with a 63x glycerol immersion objective (n=6). In situ hybridization with an RNA probe against *Nematostella Axin* was performed as described previously (*15, 17*). The anti-GFP staining intensity was measured over all ectodermal DAPI-positive nuclei starting from the deepest pharyngeal cell (relative position 0) to the cell in the middle of the aboral ectodermal domain (relative position 1) using FIJI (*47*). Briefly, to identify the ROIs, polygonal selections were drawn to separate the pharynx ectoderm and the outer ectoderm based on DAPI signal. Masks were then generated separately for the outer ectoderm and the pharynx ectoderm parts of the image using the Convert to mask and the Watershed commands. To generate the ROIs for the nuclei, particle analysis with a minimum size of 1 μm^2^ was performed. The resulting ROIs were then manually checked and sorted such that they are arranged from the relative position 0 to the relative position 1. The mean intensities in the sfGFP channel were measured for all ROIs. The relative position of each nucleus was determined as a nucleus number divided by the total number of nuclei with measured anti-GFP staining intensity in this particular embryo. *Axin* in situ staining intensity was measured in FIJI (*47*) on in situ images (n=10) along a line drawn from the deepest pharyngeal cell (relative position 0) to the cell in the middle of the aboral ectodermal domain (relative position 1).

The relative position corresponding to each measurement was determined by dividing the measurement number by the total number of measurements. In order to be able to plot the relative staining anti-GFP and *Axin* staining intensities on the same graph, for each embryo, the minimal measured staining intensity was subtracted from all intensity measurements, and then each measurement was divided by the maximum measurement.

### Live imaging

Embryos was embedded in 0.7% low-melting agarose in *Nematostella* medium (*Nematostella* medium = 16‰ artificial sea water, Red Sea Salt) in an optical bottom 35 mm Petri dish (D35-20-1.5-N, Cellvis, US) and imaged with a 20X CFI Plan Apo Lambda Objective (Nikon, Japan) using a Nikon Ti2-E/Yokogawa CSU W1 Spinning Disk Confocal Microscope. A 488 nm laser was used in conjunction with a 525/30 Emission Filter (BrightLine HC, Semrock, US) and a 25 μm pinhole size disk. Images were acquired every 5 minutes using automated imaging, over 25 Z-sections covering 120 μm depth. In the first experiment, the embryos were left to develop in 16‰ artificial sea water. In the second experiment, the embryos were developing in a 5 μM solution of the GSK3β inhibitor alsterpaullone (Sigma) from 1 hpf on. Live imaging was stopped after gastrulation was observed in the sample developing in the absence of alsterpaullone. Since embryos placed into a GSK3β inhibitor do not gastrulate, we continued the imaging of the alsterpaullone-treated embryos for an additional hour in comparison to the normal embryos. During imaging, the medium in the sample dish was continuously pumped through a tube submerged in a room temperature (∼23°C) water bath to offset heating from the microscope. This was achieved with a modified sample dish lid with two liquid connectors, and a peristaltic pump (Minipuls 3, Gilson, US). Ten embryos were imaged together in each experiment. After imaging, each Z-stack of images was converted into a maximum intensity projection.

## Author contributions

T.L. planned and performed experiments and generated the knock-in line; J.B. performed live imaging; I.N. performed *Axin* in situ hybridization; D.M. and E.G. measured and analysed the gradient data; I.A. provided access to the spinning disk confocal microscope; G.G. conceived the project, performed experiments and wrote the paper. All authors edited the paper.

## Acknowledgements

The work in Genikhovich group is funded by the Austrian Science Foundation (FWF) grants P30404 and P32705-B. The work in the Adameyko group is funded by the ERC Synergy grant “KILL-OR-DIFFERENTIATE” 856529. T.L. was a recipient of the PhD completion grant of the Vienna Doctoral School of Ecology and Evolution. D.M. is a recipient of the Lise-Meitner Fellowship M3291-B of the FWF. We thank the Core Facility for Cell Imaging and Ultrastructure Research of the University of Vienna for the access to the confocal microscope.

