## Supplementary information for "β-catenin-dependent endomesoderm specification appears to be a Bilateria-specific co-option"

#### Contents

Supplementary Figure 1

Supplementary Movie legends

Supplementary Movies 1-4

### Supplementary Figure

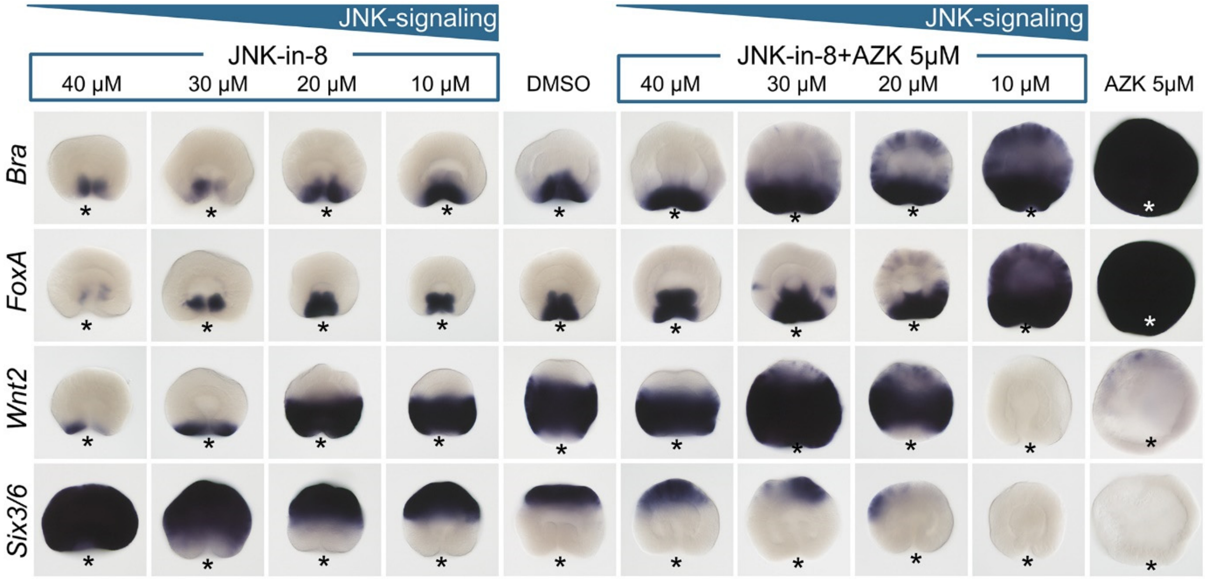


**Supplementary Figure 1. JNK-in-8 treatment dose-dependently aboralizes *Nematostella* embryos and rescues oralization caused by β-catenin stabilization with azakenpaullone (AZK).** Asterisks mark the position of the blastopore.

### Supplementary movie legends

**Supplementary movies 1 and 2. Live imaging of the sfGFP-β-catenin dynamics during development of the *Nematostella* embryo until the onset of gastrulation.** Note that nuclear sfGFP-β-catenin is visible in the interphase of every cell cycle until mid-blastula at the side opposite to where the preendodermal plate will form and start to invaginate.

**Supplementary movies 3 and 4. Live imaging of the sfGFP-β-catenin dynamics during development of the *Nematostella* embryo upon GSK3β inhibition with 5 µM alsterpaullone.** Note that nuclear sfGFP-β-catenin is localized in all nuclei throughout the embryo and keeps appearing in every cell cycle until the end of the movie, although the we filmed alsterpaullone-treated embryos for 1 hour longer than the untreated embryos shown in the Supplementary Movies 1 and 2. Also note that, as previously reported (*1, 2*), in the embryos incubated in GSK3β inhibitor from fertilization on, preendodermal plates do not form.

1. I. Niedermoser, T. Lebedeva, G. Genikhovich, Sea anemone Frizzled receptors play partially redundant roles in the oral-aboral axis patterning. *Development* **149**, dev200785 (2022).
2. L. Leclère, M. Bause, C. Sinigaglia, J. Steger, F. Rentzsch, Development of the aboral domain in Nematostella requires beta-catenin and the opposing activities of Six3/6 and Frizzled5/8. *Development* **143**, 1766-1777 (2016).
